# *Yersinia pseudotuberculosis* employs a multifaceted strategy to survive antimicrobials

**DOI:** 10.64898/2026.03.04.709318

**Authors:** Olivia Goode, Jilly Woodward, Urszula Łapińska, Jemima Onime, Audrey Farbos, Paul O’Neill, Aaron Jeffries, Christopher H. Jenkins, Isobel H. Norville, Stefano Pagliara

**Affiliations:** Living Systems Institute, University of Exeter, Stocker Road, Exeter, United Kingdom; School of Biosciences, College of Life and Environmental Sciences, University of Exeter, Stocker Road, Exeter, United Kingdom; Dstl, Porton Down, Salisbury, SP4 0JQ, United Kingdom

**Keywords:** persisters, antibiotic tolerance, Eagle effect, antibiotics, hydrogen peroxide, catalase, gene expression, *Yersinia*

## Abstract

Microbes have evolved a variety of strategies to survive exposure to naturally occurring and synthetic antimicrobials. These strategies have been investigated extensively in model bacterial organisms, whereas less is known about under explored pathogenic bacteria such as bacteria within the *Yersinia* genus. In this study we investigated the inhibitory effect and bactericidal activity of antibiotics from five different classes and of the disinfectant hydrogen peroxide against *Yersinia pseudotuberculosis*, the ancestral species from which *Yersinia pestis* and *Yersinia enterocolitica* have emerged. We found that *Y. pseudotuberculosis* is able to survive exposure to clinical antibiotics and disinfectants by employing a variety of strategies, with persisters and the Eagle effect playing a role in survival to quinolones, tolerance playing a role in survival to ceftriaxone and overexpression of catalases and peroxidases playing a role in survival to hydrogen peroxide. Our findings suggest that future research should focus on informing new, effective ways to treat infections caused by Yersinia species.

**IMPORTANCE:** Antimicrobial resistance is routinely investigated by measuring the minimum inhibitory concentration of antimicrobials needed to stop microbial growth. Here we show that the bacterial pathogen *Yersinia pseudotuberculosis* is not killed when antibiotics and disinfectants are used at these concentrations and that, in some cases, increasing antibiotic concentrations decreases their activity against this bacterium, therefore posing a potential risk to human and animal health.

## INTRODUCTION

Infectious diseases are one of the leading causes of death worldwide and antimicrobial resistance (AMR) poses a further challenge to global health with an estimated 5 million deaths associated with AMR every year (1). The misuse of antibiotics in human health and farming has accelerated the development of pathogens that are resistant to antibiotics (2–6), while the increase use of disinfectants since the coronavirus pandemic (7) raised concerns over the increase in biocide resistance (8).

Antibiotic resistance often occurs at the genotypic level: bacterial populations either encode antibiotic resistance genes or acquire genetic changes during exposure to antibiotics (9–11) or via horizontal gene transfer (12). Bacterial populations can survive antibiotics also without encoding antibiotic resistance genes. In the case of antibiotic persistence or tolerance, transient phenotypic changes in a subpopulation or in the whole bacterial population prolong survival in the presence of typically lethal concentrations of antibiotics, without the emergence of mutations (13–15). Recent evidence suggests significant increase in persisters in patients with relapsing infections (16) and a strong link between genetic resistance and persistence or tolerance (17–22), often collectively termed recalcitrance (23).

Recalcitrance to antibiotics has been investigated in depth *in vitro* (and to a lesser extent *in vivo* (23)) in bacterial model organisms, primarily *Escherichia coli* (24–27), *Salmonella enterica* (19, 28–30), *Mycobacterium tuberculosis* (31)*, Staphylococcus aureus* (32–35) and *Pseudomonas aeruginosa* (36–38). Several different processes have been identified to play a role in the recalcitrance of these organisms to antibiotics, including toxin-antitoxin systems (24, 39–41), stress response signalling molecules (42, 43), DNA damage repair systems and the SOS response (44–46), the formation of protein aggregates (47–50) and intracellular pH (15, 51).

In contrast, recalcitrance to antibiotics in other pathogens and to disinfectants have been investigated to a lesser extent (52). For example, *Yersinia* species are known to cause long-term infections, but little is known about recalcitrance to antibiotics and disinfectants in *Yersinia* (53, 54). *Yersinia* species are Gram-negative bacteria, with three species being pathogenic to humans: *Yersinia pestis, Yersinia enterocolitica, and Yersinia pseudotuberculosis*, the latter being the ancestral species from which the other two species emerged (55, 56). There are a variety of reservoirs in the environment which can serve as sources for *Yersinia* outbreaks including: mammalian and avian hosts, soil, plants, insects and contaminated water (57–63). Most cases occur in winter due to the cold tolerance of *Yersinia* which can survive and proliferate at low temperatures posing a potential food hazard (64). *Y. enterocolitica* and *Y. pseudotuberculosis* are enteric pathogens that are ingested through contaminated food, have broad animal host range (56), and can colonize both the intestine and deep tissues retaining an extracellular lifestyle (65). Progression to systemic infection is relatively rare among patients but if left untreated, patients succumb to systemic infection with an estimated mortality rate of 75%–100% (66).

Multidrug resistance has been reported for both *Y. pestis* and *Y. pseudotuberculosis* and it has been linked to the acquisition of plasmids containing multiple resistance genes (67, 68). Moreover, around 10% of a *Y. pseudotuberculosis* population infecting mice spleen survived a single inhibitory dose of doxycycline (53). This subpopulation was primarily constituted of bacteria at the periphery of microcolonies that responded to host-produced nitric oxide and other stresses by expressing the detoxifying gene, *hmp* (69). This subpopulation may have persisted for days and resumed growth between 48 and 72 h posttreatment when the doxycycline concentration declined, ultimately leading to mice death (53). At a population level, upregulation of the genes encoding the outer membrane protein OmpF and the outer membrane lipoprotein OsmB was recorded in response to inhibitory concentrations of doxycycline both *in vitro* and *in vivo*, together with downregulation of the genes encoding the sulfur transfer complex subunit TusB and the cytotoxic necrotizing factor CnfY (54). OmpF seemed to perform the counterintuitive function of promoting diffusion of doxycycline out of the cell, whereas overexpression of *tusB* increased sensitivity to doxycycline (54). It has also been shown that the induction of *Yersinia* type-III secretion system leads to growth inhibition and reduced susceptibility to the antibiotic gentamicin *in vitro*, and that *Y. pseudotuberculosis* bacteria surviving gentamicin in a mouse model also express low levels of *rpsJ*, which encodes the 30S ribosomal protein S10 (70).

These studies demonstrated that *Y. pseudotuberculosis* can display persistence or tolerance to gentamicin or doxycycline treatment. However, it remains unclear whether *Y. pseudotuberculosis* can display persistence or tolerance to high concentrations of a wider range of antibiotics and disinfectants that are routinely employed for antibiotic therapy or decontamination purposes. Here we measure and compare susceptibility, persistence and tolerance of *Y. pseudotuberculosis* to supra-inhibitory concentrations of antibiotics from five different classes recommended as first-line antibiotics by current guidelines from Centers for Disease Control and Prevention (71), and of hydrogen peroxide, which is used as a topical antiseptic and as an effective disinfectant (72–74). We further measure and compare the survival of the *Y. pseudotuberculosis* population after a 24 hour exposure to supra-inhibitory concentrations of antibiotics or of hydrogen peroxide. We observed extensive survival of *Y. pseudotuberculosis* after 24 hour exposure to high concentrations of hydrogen peroxide and therefore carried out genome-wide comparative transcriptome analysis of *Y. pseudotuberculosis* to identify changes in gene expression in response to the exposure to hydrogen peroxide. Collectively, our *in vitro* data demonstrate that *Y. pseudotuberculosis* is difficult to kill via either antibiotics or disinfectants and suggest that future research should focus on informing new, effective ways to treat *Y. pseudotuberculosis*.

## RESULTS

### Minimum inhibitory concentrations of antimicrobials are not sufficient to kill *Yersinia pseudotuberculosis*

Using the broth microdilution assay (21), we investigated the susceptibility of *Y. pseudotuberculosis* to antimicrobial compounds while the bacterium was in two distinct phases of growth, namely exponential phase or stationary phase, i.e. after 6 h or 17 h of culture in Lysogeny broth (LB) medium at 37°C with shaking at 200 rpm, respectively. Indeed, *Y. pseudotuberculosis* remained in the lag phase for the first 4 hours of exposure to fresh LB, grew in exponential phase between 4 and 14 hours and remained in stationary phase between 14 and 24 hours (Supplementary Figure 1), in accordance with previous reports (64, 75, 76).

We found that *Y. pseudotuberculosis* was equally susceptible when treated with antibiotics during exponential or stationary phase with the lowest minimum inhibitory concentration (MIC) value recorded for ceftriaxone (0.25 µg ml^-1^) and the highest MIC value recorded for gentamicin and chloramphenicol (4 µg ml^-1^). *Y. pseudotuberculosis* was instead slightly more susceptible to hydrogen peroxide when treated during exponential phase compared to stationary phase with MIC values of 1.25 mM and 2.5 mM, respectively (Table 1). Consistently, using colony forming unity (CFU) assays (77), we found that the minimum bactericidal concentration (MBC) value of hydrogen peroxide was 8 mM against exponential phase *Y. pseudotuberculosis* and 16 mM against stationary phase *Y. pseudotuberculosis*. Similarly, the MBC value of ceftriaxone was slightly higher when *Y. pseudotuberculosis* was treated in stationary phase compared to exponential phase (8 µg ml^-1^ and 4 µg ml^-1^, respectively). The lowest MBC value was recorded for ciprofloxacin and levofloxacin (2 µg ml^-1^), whereas the highest MBC values were recorded for doxycycline and chloramphenicol (> 64 µg ml^-1^) that are two bacteriostatic antibiotics (Table 1), although it has recently been suggested that doxycycline can also have bactericidal activity against *Y. pseudotuberculosis* (54). A relatively large discrepancy between the MIC and MBC values (> 4-fold) was measured for doxycycline, gentamicin, ceftriaxone and chloramphenicol, whereas a relatively small discrepancy (< 4-fold) was measured for the two fluoroquinolones.

**Table 1.** *Y. pseudotuberculosis* susceptibility to antimicrobials. MIC and MBC values measured for seven different antimicrobials against *Y. pseudotuberculosis* either in exponential or stationary phase. Values are reported in µg ml^-1^ for the six antibiotics tested and in mM for hydrogen peroxide. Values were obtained as the mean of measurements carried out in biological triplicates each consisting of technical triplicates.

| | MIC values ($\mu\text{g ml}^{-1}$) | | MBC values ($\mu\text{g ml}^{-1}$) | |
| --- | --- | --- | --- | --- |
|  | Exp. phase | Stat. phase | Exp. phase | Stat. phase |
| Doxycycline | 1 | 1 | > 64 | > 64 |
| Levofloxacin | 1 | 1 | 2 | 2 |
| Ciprofloxacin | 0.5 | 0.5 | 2 | 2 |
| Gentamicin | 4 | 4 | 32 | 32 |
| Ceftriaxone | 0.25 | 0.25 | 4 | 8 |
| Chloramphenicol | 4 | 4 | > 64 | > 64 |
| Hydrogen peroxide | 1.25 | 2.5 | 8 | 16 |

### *Y. pseudotuberculosis* recalcitrance to antibiotics and hydrogen peroxide

Next, we set out to investigate both antimicrobial tolerance at the population level and the presence and levels of persisters to antimicrobials within clonal populations of *Y. pseudotuberculosis* in the stationary phase of growth by using supra-MIC antimicrobial concentrations and CFU assays.

We found that the aminoglycoside gentamicin displayed slow killing when used at 1× its MIC value (i.e. 4 µg ml^-1^) with a minimum duration of killing (MDK_99_) longer than 5 h (Figure 1A), suggesting *Y. pseudotuberculosis* tolerance to gentamicin at the population level and at this concentration. However, when this antibiotic was used at 10× or 25× its MIC value (i.e. 40 µg ml^-1^ thus slightly above gentamicin’s MBC value and 100 µg ml^-1^, respectively), the MDK_99_ was less than 1 h with a complete killing of the *Y. pseudotuberculosis* population and no detectable survivors (Figure 1A).

**Figure 1.**
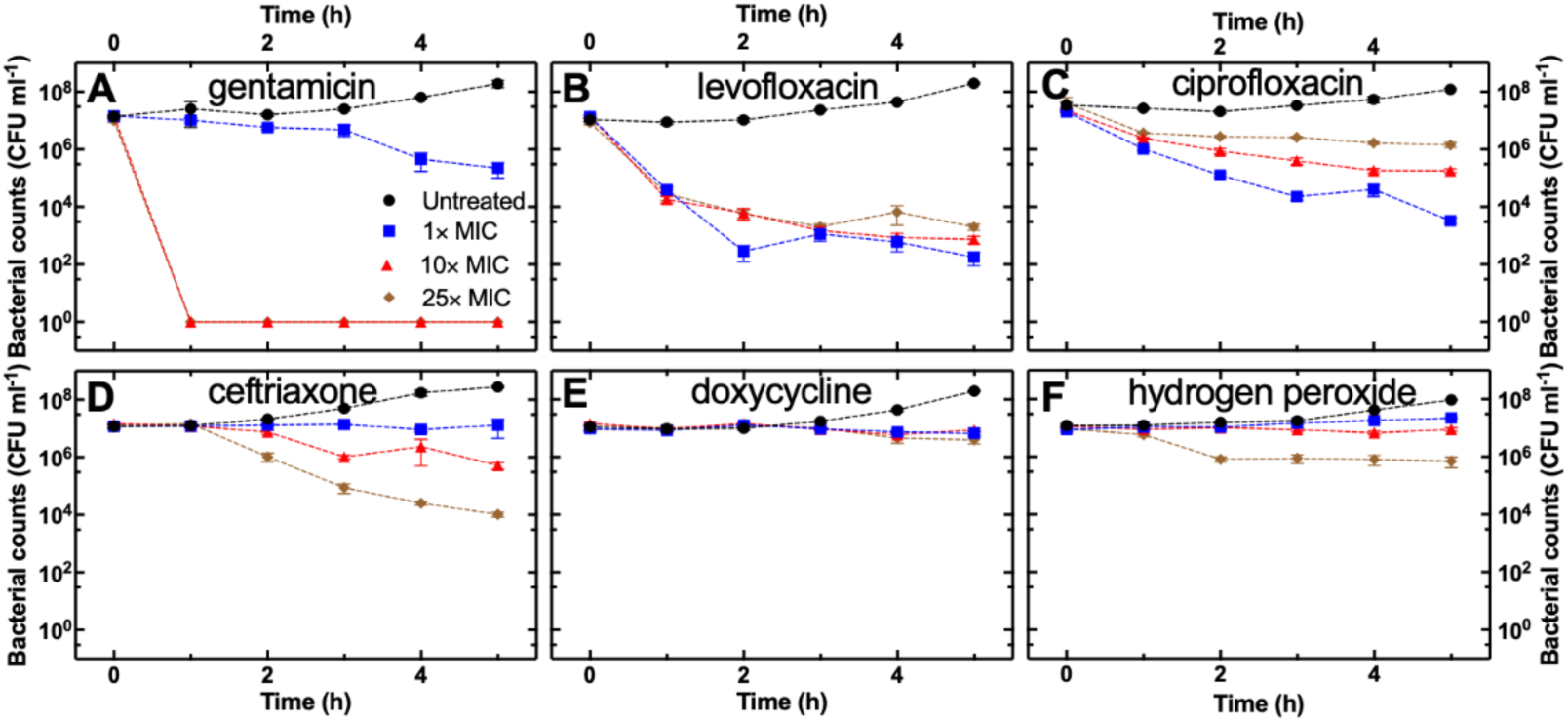
*Y. pseudotuberculosis* recalcitrance to antimicrobials. Time-dependent killing of *Y. pseudotuberculosis* by a) gentamicin, b) levofloxacin, c) ciprofloxacin, d) ceftriaxone, e) doxycycline or f) hydrogen peroxide used at their MIC value (blue squares), 10× (red triangles) or 25× their MIC value (brown diamonds). Corresponding data for untreated control samples are reported with black circles in each graph. Symbols and error bars are the mean and standard error of biological triplicate each consisting of technical triplicate.

The third-generation fluoroquinolone levofloxacin displayed potent bactericidal activity with an MDK_99_ of less than 1 h already when this compound was used at 1× its MIC value (i.e. 1 µg ml^-1^, Figure 1B), suggesting that *Y. pseudotuberculosis* is not tolerant to levofloxacin at the population level. After this initial rapid killing, the *Y. pseudotuberculosis* population did not further decrease from 2 h onwards, the overall population dynamics following biphasic killing which indicates the presence of persisters (14). The *Y. pseudotuberculosis* persister fraction was 10^-5^ (i.e. one persister every hundred thousand cells in the initial bacterial population) when levofloxacin was used at 1× its MIC value; surprisingly, increasing levofloxacin concentration to 10× or 25× its MIC value (i.e. 10 µg ml^-1^ and 25 µg ml^-1^, respectively) increased the persister fraction to 5×10^-5^ and 10^-4^, respectively, after 5 h treatment (Figure 1B). We found a similar trend also when we employed increasing concentrations of the second-generation fluoroquinolone ciprofloxacin, with the MDK_99_ increasing from 2 h to 4 h and to more than 5 h and the *Y. pseudotuberculosis* persister fraction increasing from 10^-4^ to 10^-2^ and to 10^-1^ when ciprofloxacin was used at 1×, 10× or 25× its MIC value (i.e. 0.5 µg ml^-1^, 5 µg ml^-1^, and 12.5 µg ml^-1^, respectively, Figure 1C). We also noted a slower more gradual killing of *Y. pseudotuberculosis* by ciprofloxacin compared to levofloxacin which is reflected in the larger MDK_99_ values and the less pronounced biphasic killing trends (compare Figure 1B and 1C), suggesting *Y. pseudotuberculosis* tolerance to high concentrations of ciprofloxacin.

The third-generation cephalosporin ceftriaxone did not display bactericidal activity against *Y. pseudotuberculosis* when used at its MIC value (i.e. 0.25 µg ml^-1^) in accordance with the corresponding MBC data above. The MDK_99_ decreased from more than 5 h to 3 h and the *Y. pseudotuberculosis* survival fraction decreased from 4×10^-2^ to 10^-3^ when the ceftriaxone concentration was increased from 10× to 25× its MIC value (i.e. 2.5 µg ml^-1^ and 6.25 µg ml^-1^, respectively, Figure 1D). Killing of *Y. pseudotuberculosis* was slow and gradual, suggesting tolerance to ceftriaxone at the population level and it is also worth noting that no killing was recorded during the first hour of treatment with ceftriaxone (Figure 1D). As expected, doxycycline being a bacteriostatic antibiotic, did not display bactericidal activity against *Y. pseudotuberculosis* at any of the concentrations tested (Figure 1E).

Finally, the disinfectant hydrogen peroxide displayed bactericidal activity against *Y. pseudotuberculosis* only when employed at 25× its MIC value (i.e. 62.5 mM) with a MDK_99_ longer than 5 h and a survival fraction of 10^-1^ (Figure 1F), suggesting tolerance to hydrogen peroxide at the population level. Taken together these data suggest that among the antimicrobials tested, fluoroquinolones provide the strongest bactericidal activity against *Y. pseudotuberculosis* at the lowest concentrations; however, the *Y. pseudotuberculosis* survival fraction depends strongly on the fluoroquinolone concentration which might be difficult to optimise *in vivo*.

### *Y. pseudotuberculosis* growth after prolonged exposure to antimicrobials

Next, we carried out CFU assays after exposing *Y. pseudotuberculosis* to each of the antimicrobials above for 24 h. Consistent with the data above, we found that when used at its MIC value gentamicin caused a 3-log reduction in the number of culturable *Y. pseudotuberculosis* cells compared with untreated controls; whereas we did not detect any culturable *Y. pseudotuberculosis* cells when gentamicin was used at 10× or 25× its MIC value (Figure 2A). Similarly, we did not detect any culturable *Y. pseudotuberculosis* cells after a 24 h exposure to levofloxacin at 1×, 10× or 25× its MIC value (Figure 2B). We measured a 4-log, 6-log and 5-log reduction in the number of culturable *Y. pseudotuberculosis* cells compared with untreated controls when ciprofloxacin was used at 1×, 10× or 25× its MIC value, respectively (Figure 2C). We measured a 5-log and 7-log reduction in the number of culturable *Y. pseudotuberculosis* cells compared with untreated controls when ceftriaxone was used at 1× or 10× its MIC, respectively, whereas we did not detect any culturable *Y. pseudotuberculosis* cells when ceftriaxone was used at 25× its MIC value (Figure 2D). We measured a 3-log, 4-log and 4-log reduction in the number of culturable *Y. pseudotuberculosis* cells compared with untreated controls when doxycycline was used at 1×, 10× or 25× its MIC value (Figure 2E). Prolonged exposure to hydrogen peroxide yielded a 0.5-log, 1-log and 2-log reduction in the number of culturable *Y. pseudotuberculosis* cells compared with untreated controls when used at 1×, 10× or 25× its MIC value, respectively (Figure 2F). Together with the recalcitrance data above, these data suggest that levofloxacin is the most effective antimicrobial against *Y. pseudotuberculosis in vitro* and raise concerns about the use of other antimicrobials, particularly hydrogen peroxide, that is often employed as a disinfectant or a decontamination agent.

**Figure 2.**
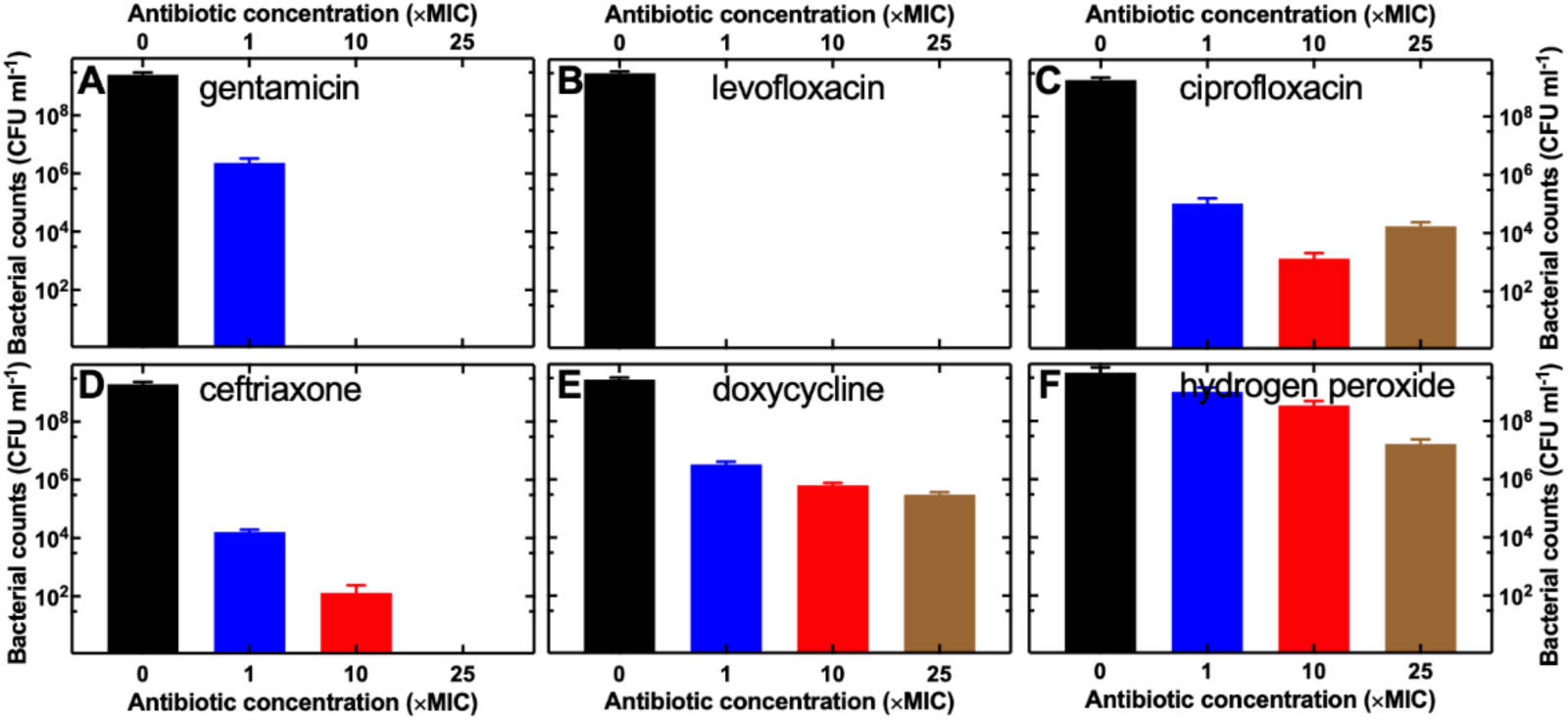
*Y. pseudotuberculosis* growth after exposure to supra-MIC concentrations of antimicrobials. Counts of *Y. pseudotuberculosis* cells after 24 h exposure to a) gentamicin, b) levofloxacin, c) ciprofloxacin, d) ceftriaxone, e) doxycycline or f) hydrogen peroxide used at 1×, 10×, or 25× their MIC value. Bars and error bars are the mean and standard error of biological triplicate each consisting of technical triplicate. Each set of experiments with a different antimicrobial included untreated control experiments without adding the antimicrobial, *Y. pseudotuberculosis* growth in these experiments is reported by the black bars (i.e. 0× the MIC value).

### *Y. pseudotuberculosis* is more susceptible to a second exposure to hydrogen peroxide

We further investigated *Y. pseudotuberculosis* survival to hydrogen peroxide exposure and found a transition from *Y. pseudotuberculosis* population tolerance to subpopulation persistence when the concentration of hydrogen peroxide was increased from 25× to 100× its MIC value, with an MDK_99_ decreasing from more than 5 h to less than 1 h (Figure 3A). The *Y. pseudotuberculosis* persister fraction after exposure to hydrogen peroxide at 100× its MIC value was 10^-5^; this low number of cells was sufficient for *Y. pseudotuberculosis* regrowth to 0.5 million cells ml^-1^ after 24 h in the presence of hydrogen peroxide (Figure 3A). *Y. pseudotuberculosis* was fully killed only after exposure to hydrogen peroxide used at 400× its MIC value (i.e. 1M) with a MDK_99_ value of less than 1 h and no detectable persisters or regrowth after 24 h (Figure 3A).

**Figure 3.**
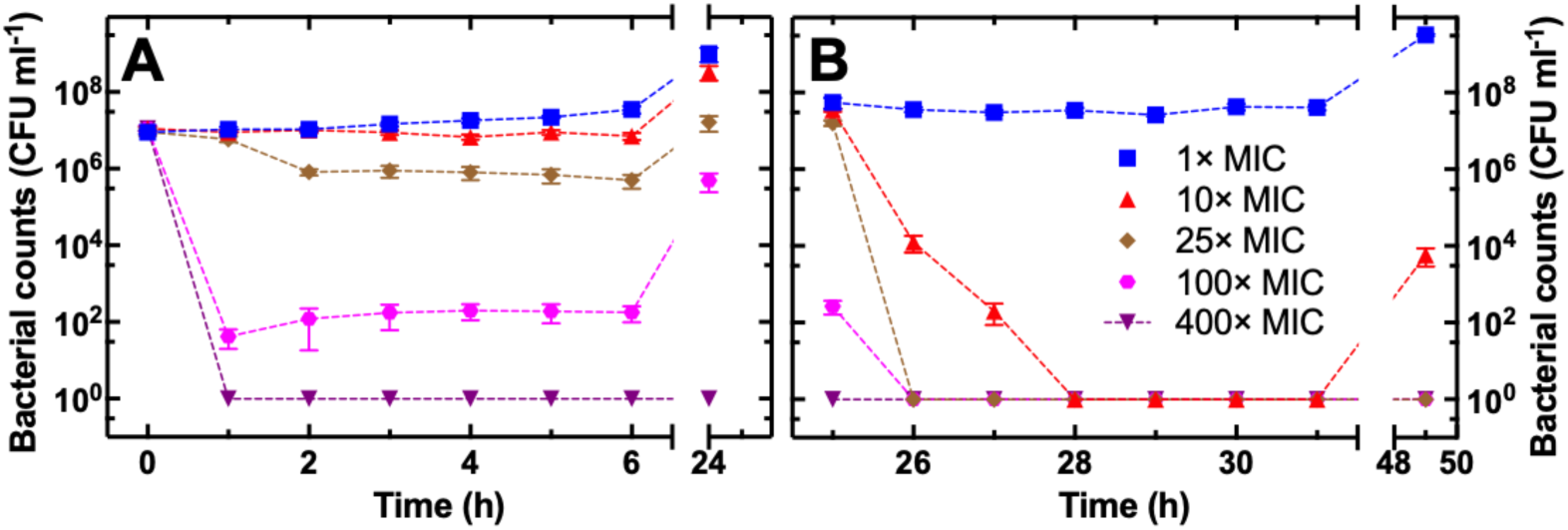
*Y. pseudotuberculosis* is more susceptible to a second exposure to hydrogen peroxide. Time-dependent killing of *Y. pseudotuberculosis* by a) a first and b) a second exposure to hydrogen peroxide used at its MIC value (blue squares), 10× (red triangles), 25× (brown diamonds), 100× (magenta hexagons), or 400× its MIC value (purple downward triangles). Symbols and error bars are the mean and standard error of biological triplicate each consisting of technical triplicate.

We then collected the *Y. pseudotuberculosis* cells that had survived different concentrations of hydrogen peroxide (i.e. 24 h time point in Figure 3A), washed them in PBS to remove any residual hydrogen peroxide, resuspended them in 100 ml of LB and exposed them a second time to the same concentration of hydrogen peroxide for another 24 h.

We found that *Y. pseudotuberculosis* was not killed by a second exposure to hydrogen peroxide at its MIC value, similarly to what we observed during the first exposure; however, when hydrogen peroxide was used at 10× its MIC value, *Y. pseudotuberculosis* regrowth after the second exposure was significantly lower compared to the first exposure (i.e. 10^3^ cells ml^-1^ vs 10^8^ cells ml^-1^, respectively, p-value < 0.0001 according to unpaired t-test), and we did not detect any persisters or regrowth during the second exposure to hydrogen peroxide at 25×, 100× or 400× its MIC value (Figure 3B).

Taken together these data demonstrate that i) *Y. pseudotuberculosis* survival and regrowth after the first exposure to hydrogen peroxide is not due to stable genetic mechanisms that can be passed on to the *Y. pseudotuberculosis* progeny; ii) *Y. pseudotuberculosis* is more susceptible to a second exposure to hydrogen peroxide and therefore repeated applications of this antimicrobial agent can lead to *Y. pseudotuberculosis* complete killing provided that its concentration is sufficiently high, i.e. at least 25× its MIC value or 62.5 mM under our *in vitro* experimental conditions. In contrast, complete killing via a single exposure can be achieved only by using a very high concentration of hydrogen peroxide, i.e. 400× its MIC value or 1 M.

### Gene expression changes triggered by exposure to hydrogen peroxide

Next, we employed an unbiased approach to examine changes in gene expression by performing genome-wide comparative transcriptome analysis (78, 79) on *Y. pseudotuberculosis* after 6 h or 24 h of culture in LB or after 6 h or 24 h exposure to hydrogen peroxide at a concentration of 2.5 mM or 62.5 mM (i.e. 1× or 25× its MIC value). Samples were prepared for sequencing using our previous protocols for RNA-seq (78, 79) and the readings aligned to the *Y. pseudotuberculosis* genome (https://www.ncbi.nlm.nih.gov/nuccore/BX936398, see S1 File for RNA transcript counts for each replicate and S2 File for differential expression of individual genes in each condition compared to the corresponding untreated control experiments). Principal component analysis revealed major divergences across different culturing conditions and high within group reproducibility, i.e. transcriptome replicates from each condition clustered together (Figure S2).

As expected, 6 h exposure to hydrogen peroxide at a concentration of 2.5 mM elicited weaker differential gene expression compared to a concentration of 62.5 mM (Figure 4A-4B). Notably, the genes encoding the catalase-peroxidase and the catalase enzymes KatG and KatA, were upregulated 6.8 and 5.8 log_2_ fold after 6 h exposure of *Y. pseudotuberculosis* to 62.5 mM hydrogen peroxide; *cybB* and *cybC*, encoding the cytochromes b561 and b562, were upregulated 7.5 and 6.7 log_2_ fold, the genes encoding the acid-activated periplasmic chaperone HdeB, the outer membrane protein YcoB and the reductase TrxC were upregulated 5.8, 4.8 and 5.5 log_2_ fold, respectively (Figure 4B and S2 File). On the other hand, the genes encoding the nitrate reductase complex NapAB, the cytochrome NapC, the chaperone NapD and the ferredoxin-type protein NapF were downregulated 4.0, 2.4, 1.6, 5.7 and 4.9 log_2_ fold, respectively; the genes encoding the nitrite reductase subunit NirB and NirD and the nitrite transporter NirC were downregulated 4.6, 4.8 and 3.1 log_2_ fold, respectively; *cydA* and *cydB*, encoding the cytochrome ubiquinol oxidase subunits I and II, were downregulated 2.6 and 1.8 log_2_ fold, respectively; the gene encoding the 5-oxoprolinase subunit PxpB was downregulated 2.8 fold (Figure 4B and S2 File).

**Figure 4.**
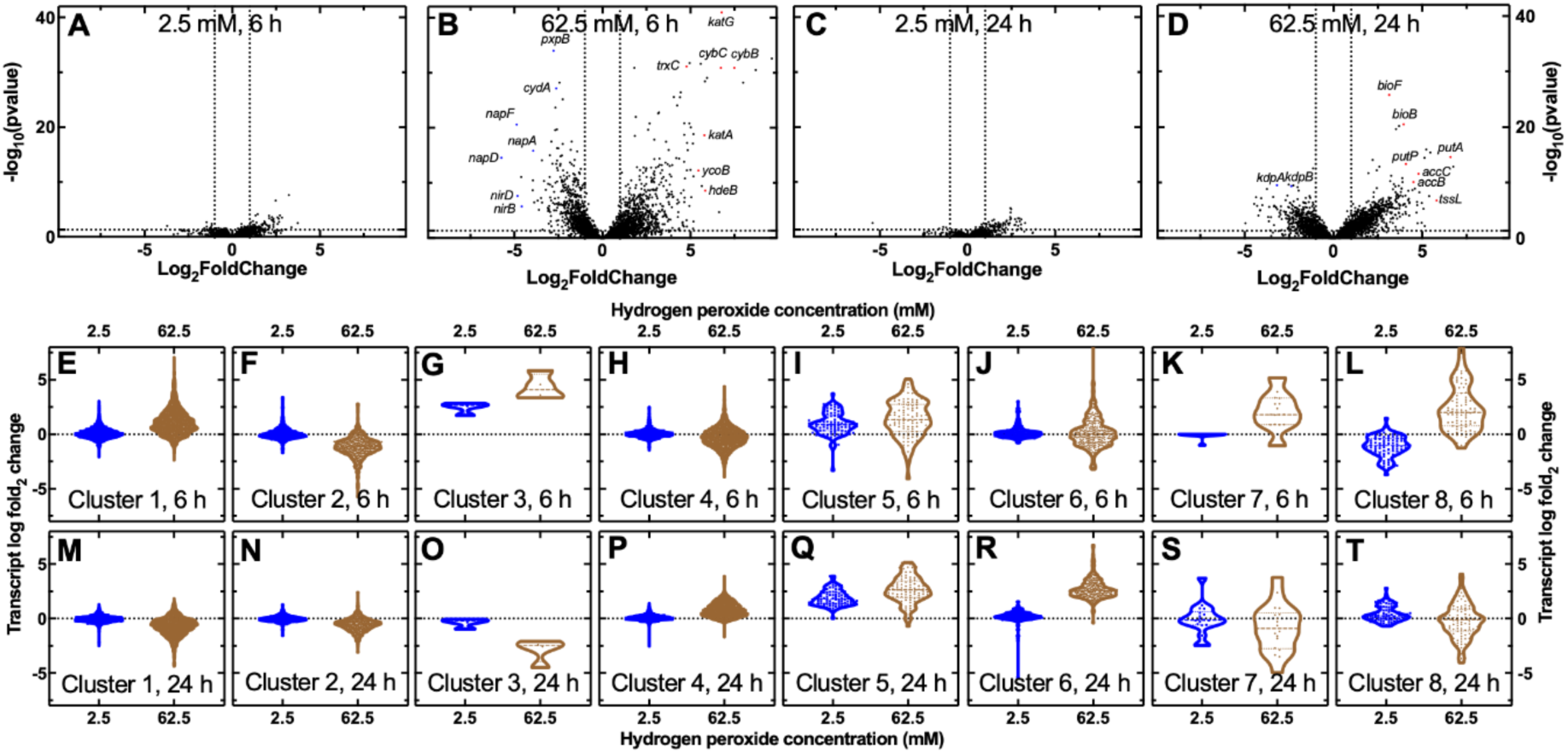
*Y. pseudotuberculosis* transcriptome rewiring in response to hydrogen peroxide. Volcano plot of whole-genome differential gene expression in *Y. pseudotuberculosis* after (a) 6 h exposure to 2.5 mM or (b) 62.5 mM hydrogen peroxide or (c) 24 h exposure to 2.5 mM or (d) 62.5 mM hydrogen peroxide compared to gene expression in corresponding untreated control experiments. Each dot represents the average log_2_ fold change in transcript reads of each gene and the corresponding adjusted p-value. Horizontal and vertical dotted lines indicate an adjusted p-value of 0.05 and a log_2_ fold change of 2, respectively. Strongly upregulated or downregulated genes of interest are highlighted in red and blue respectively. Cluster analysis of differential gene expression in *Y. pseudotuberculosis* after e)-l) 6 h or m)-t) 24 h exposure to either 2.5 mM (blue dots and graphs) or 62.5 mM (brown dots and graphs) of hydrogen peroxide compared to untreated *Y. pseudotuberculosis* controls. Each dot represents the log_2_ fold change in transcript reads for a single gene. Dotted lines indicate a log_2_ fold change of zero. The lists of genes belonging to each cluster are reported in S2 File. Each measurement was carried out in biological triplicate and the values reported are the mean of these measurements (expressed as log_2_ fold change compared to untreated controls). Transcript read counts for the untreated controls as well as the error and p-value associated to each comparison are reported in S2 File. RNA transcript counts for each replicate are reported in S1 File.

After 24 h exposure to hydrogen peroxide at a concentration of 62.5 mM a different set of genes was strongly differentially regulated: the genes encoding the proline dehydrogenase PutA and the sodium/proline symporter PutP, the type IV secretion system protein TssL, the acetyl-CoA carboxylase subunits AccB and AccC and the biotin synthases BioB and BioF were upregulated 6.6, 4.1, 5.8, 4.6, 4.8, 4.0 and 3.2 log_2_ fold, respectively; the genes encoding the potassium-transporting ATPase subunits KdpA and KdpB were downregulated 3.2 and 2.4 log_2_ fold, respectively (Figure 4D).

Next, in order to identify biological processes that were differentially regulated in *Y. pseudotuberculosis* due to exposure to hydrogen peroxide, we performed cluster analysis (15, 80) of our transcriptomic data and identified 8 distinct patterns of gene regulation.

Genes in clusters 1, 3, 7 and 8 displayed an initial upregulation after 6 h exposure to 62.5 mM hydrogen peroxide compared to exposure to 2.5 mM hydrogen peroxide or untreated controls (Figure 4E, 4G, 4K, 4L); however, these genes were downregulated after 24 h exposure to 62.5 mM hydrogen peroxide (Figure 4M, 4O, 4S, 4T). Cluster 1 contained 1582 genes and by using gene ontology analysis (35) we determined that this cluster was enriched for genes in cell adhesion, membrane transport, protein folding and hydrolase activity processes (see S3-S10 Files for all biological processes enriched in each cluster). Clusters 3 and 7 contained only 4 and 18 genes, respectively. Cluster 8 contained 87 genes and was enriched for genes in response to oxidative stress (including *katA* and *katG*), cell redox homeostasis, electron transfer activity (including *cybB* and *cybC*) and tricarboxylic acid cycle processes (including *aceA*, *aceB* and *aceK*).

Cluster 2 contained 511 genes that were mostly downregulated both after 6 h and 24 h exposure to 62.5 mM of hydrogen peroxide and was enriched for sugar phosphotransferase system, anaerobic respiration, DNA-templated transcription and DNA-binding transcription factor activity processes.

Clusters 4 and 6 contained 1449 and 276 genes that were upregulated only after 24 h exposure to 62.5 mM of hydrogen peroxide. This cluster was enriched for genes in cell division, cell wall organisation, ATP binding and hydrolysis, ribosome and chromosome segregation processes. The entire NADH-quinone oxidoreductase pathway (in range 2-3.5 log_2_ fold), genes encoding an erythritol/L-threitol dehydrogenase and a D-threitol dehydrogenase (5 log_2_ fold) were upregulated after 24 h exposure to 62.5 mM of hydrogen peroxide. We also noted the strong upregulation (9 log_2_ fold) of a gene encoding an aldehyde dehydrogenase after 6 h (but not after 24 h) exposure to 62.5 mM of hydrogen peroxide, consistent with the regulation of genes in cluster 8 above.

Finally, cluster 5 contained 95 genes that were mostly upregulated both after 6 h and 24 h exposure to 62.5 mM of hydrogen peroxide. This cluster was enriched for genes in transmembrane transporter activity, cytolysis and phage related processes. This cluster included genes encoding a gluconate 5-hydrogenase (3.5 log_2_ fold change at 24 h), a NADPH-dependent FMN reductase (*ssuE*, 3 log_2_ fold change at 24 h) and taurine dioxygenase (*tauD*, 3 log_2_ fold change at 24 h)

## Discussion

*Y. pseudotuberculosis* is a model organism for *Y. pestis,* which is the causative agent of plague and is on the WHO list of high risk organisms that could cause the next pandemic (81). Indeed, Madagascar witnessed a major plague outbreak in 2017 with at least one *Y. pestis* strain displaying antimicrobial resistance (82). Here we set out to investigate persistence and tolerance to antibiotics and disinfectants in *Y. pseudotuberculosis* considering that recent evidence suggests significant increase in persisters in patients with relapsing infections (16) and a strong link between genetic resistance and persistence or tolerance in other pathogenic bacteria (17–22).

The minimum inhibitory concentration values we recorded for doxycycline, levofloxacin, ciprofloxacin and gentamicin against *Y. pseudotuberculosis* are in line with the range of values in previous reports (53, 54, 83–85). These MIC values are above the breakpoint values recommended by EUCAST for the use of these antibiotics against Enterobacterales. Moreover, although relatively high, these antibiotic concentrations are not sufficient to kill *Y. pseudotuberculosis*, the minimum bactericidal concentration values being between 2-fold and 64-fold larger than the corresponding minimum inhibitory concentration values measured for each antibiotic. The scarce efficacy of these clinical antibiotics is further exacerbated by *Y. pseudotuberculosis* also displaying either persistence or tolerance to levofloxacin, ciprofloxacin and ceftriaxone which leads to the regrowth of *Y. pseudotuberculosis* after 24 h exposure to these antimicrobials. Only the use of gentamicin at a minimal concentration of 40 µg ml^-1^ or levofloxacin at 1 µg ml^-1^ led to *Y. pseudotuberculosis* complete killing after 24 h treatment in our *in vitro* conditions.

These data demonstrate that *Y. pseudotuberculosis* is difficult to treat with antibiotics under *in vitro* conditions and are in line with recent reports demonstrating that *Yersinia pseudotuberculosis* can survive doxycycline treatment within host mouse tissues (53, 54). The strong efficacy we recorded for levofloxacin *in vitro* is also in line with previous evidence of good efficacy of levofloxacin and poor efficacy of doxycycline against pneumonic plague in a nonhumane primate model (86).

Extensive investigations have been carried out on persistence and tolerance to antibiotics, primarily using bacterial model organisms, whereas little is known about persistence and tolerance in *Yersinia* species. Persisters to meropenem and to multiple antibiotics, including ciprofloxacin, were found in attenuated *Y. pestis* (87) and in *Y. ruckerii* (88), respectively, whereas a subpopulation of *Y. enterocolitica* cells was able to survive whole blood complement (89). The new data we obtained here for *Y. pseudotuberculosis* are in line with some previous report of persisters in different species. For example, a biphasic kill curve characteristic of persister presence was recorded when levofloxacin was used at a concentration of 5 µg ml^-1^ against *E. coli* (46), in line with our data showing *Y. pseudotuberculosis* persisters to levofloxacin at a concentrations of 10 µg ml^-1^. *Y. ruckeri* persisters to ciprofloxacin were observed when this antibiotic was used at 10 µg ml^-1^ (88) and we also found *Y. pseudotuberculosis* persisters when using ciprofloxacin at 12.5 µg ml^-1^.

Notably, while for gentamicin, ceftriaxone and hydrogen peroxide bactericidal efficacy increased with antibiotic concentration as expected (90–92), the opposite was true for the two fluoroquinolones tested, ciprofloxacin and levofloxacin. This paradoxical effect, known as the Eagle effect, was first identified in 1948 when it was observed that there was an optimal penicillin concentration beyond which the efficacy of this antibiotic in killing several bacteria species, including *Streptococci*, sharply decreased (93). Investigations of the Eagle effect have been carried out in Gram-positive and Gram-negative bacteria, mycobacteria, and fungi using a diverse range of antimicrobials; however, its underpinning mechanisms are not well understood, partly due to the fact that the Eagle effect seems to be both antibiotic- and strain-dependent (94).

The Eagle effect has been observed for ciprofloxacin against *E. coli* (95), *Mycobacterium smegmatis* (96), *S. aureus*, *S. pneumoniae* and *P. aeruginosa* (97), whereas here we report for the first time this effect for levofloxacin, and for ciprofloxacin against a *Yersinia* species. Our data showing that ciprofloxacin efficacy against *Y. pseudotuberculosis* decreases at concentrations that are from 10× to 25× the MIC value (i.e. 5-12.5 µg ml^-1^) are in line with previous data: ciprofloxacin efficacy against *S. aureus*, *S. pneumoniae* and *P. aeruginosa* decreased at concentrations above 35× the MIC value (i.e. 14 µg ml^-1^) when bactericidal activity was measured in serum ultrafiltrate (97). However, it is worth reiterating that the Eagle effect appears to be strain-dependent: for example we and others have not previously observed the Eagle effect for ciprofloxacin against *E. coli* (21) or *S. aureus* (92). Postulated mechanisms behind the Eagle effect induced by quinolone antibiotics are related to suppression of reactive oxygen species accumulation at high quinolone concentrations (98). Future investigations should test whether this mechanism plays a role in the Eagle effect we observed when using fluoroquinolones against *Y. pseudotuberculosis*.

Considering that animal models support that the Eagle effect observed *in vitro* translates to *in vivo* consequences, including in human patients (94), further investigation of this effect in *Yersinia* species within host mouse tissues (53, 54) could lead to clinical benefits, where low antibiotic exposures may expedite clearance of infection. Moreover, recent evidence suggests an increase in ceftriaxone persisters in relapsed *E. coli* infection (16) and that *S. typhimurium* persisters into the gut promote resistance-plasmid transfer between other gut Enterobacteriaceae (19). Therefore, it would be valuable to investigate whether high persister isolates are also present in relapsed infecting strains of *Yersinia* species and the minimum infectious dose of persisters that causes an infection to relapse *in vivo*. It would also be valuable to investigate the mechanisms underpinning *Yersinia* persisters to antibiotics considering that mechanistic insights are so far limited to doxycycline only (53, 54).

We found that *Y. pseudotuberculosis* was difficult to eradicate also by using hydrogen peroxide, which is employed as a topical antiseptic and as an effective disinfectant (72–74). Previous reports suggested that hydrogen peroxide can be either bacteriostatic or bactericidal depending on the concentration employed (76, 99, 100). In line with these observations, we show that hydrogen peroxide starts to display some bactericidal activity against *Y. pseudotuberculosis* when used at concentrations higher than 10× its MIC value. We found a biphasic kill curve characteristic of persister presence when we used hydrogen peroxide at 100× its MIC value, whereas recent reports identified *E. coli* persistence to benzalkonium chloride used at 3× its MIC value, but no persisters to hydrogen peroxide (101, 102). Complete killing of *Y. pseudotuberculosis* was obtained only when hydrogen peroxide was used at 400× its MIC value, that is 1 M concentration or via two 24 h exposures to hydrogen peroxide at 25× its MIC value, suggesting a possible alternative to the use of highly concentrated hydrogen peroxide solutions.

Bacteria employ catalases and peroxidases to convert hydrogen peroxide to non-toxic oxygen and water, thus reducing intracellular concentrations of hydrogen peroxide (103). Catalase deletion mutants of planktonic *Salmonella enterica* serovar Typhimurium have displayed decreased MIC values and reduced tolerance in biofilms to hydrogen peroxide (104). Catalase mutants in *E. coli* accumulate enough hydrogen peroxide to cause substantial damage to DNA and display growth and viability defects (105). Furthermore, RNA transcriptome analysis of *Y. pseudotuberculosis* after five minutes of inhibitory concentrations of hydrogen peroxide revealed enrichment of genes involved in combatting oxidative stress, including OxyR regulon members and downregulation of metabolic enzymes (100). The activation of OxyR in response to hydrogen peroxide was also identified in *E. coli* after hydrogen peroxide exposure (106).

Using RNA sequencing we found that *Y. pseudotuberculosis* strongly upregulates the genes encoding the catalase-peroxidase and the catalase enzymes KatG and KatA after 6 h exposure to a supra-inhibitory and bactericidal concentration of hydrogen peroxide. More broadly, the oxidative stress, cell redox homeostasis, electron transfer activity and tricarboxylic acid cycle biological processes were upregulated at this time point. However, these genes and biological processes were downregulated after 24 h incubation in hydrogen peroxide, suggesting that by using these processes *Y. pseudotuberculosis* had decreased both the intracellular and extracellular concentration of hydrogen peroxide by this time point allowing growth to restart. Interestingly, the genes and processes above were not upregulated after exposure to an inhibitory but not bactericidal concentration of hydrogen peroxide, suggesting that *Y. pseudotuberculosis* can tolerate this concentration of hydrogen peroxide by being in a non-growing state and with a minor transcriptome rewiring.

Future investigations should directly measure how catalases and peroxidases, produced by *Yersinia* species, impact the extracellular concentration of hydrogen peroxide, whereas catalase negative mutants could be used to determine the impact of these enzymes on the MIC and MBC values of hydrogen peroxide against *Yersinia*. It would also be important to investigate cross tolerance and persistence by exposing *Yersinia* to both hydrogen peroxide and antibiotics, considering that exposure to hydrogen peroxide has been reported to increase the percentage of *E. coli* persisters to antibiotics (101, 107, 108)

In summary, we show that *Y. pseudotuberculosis* is able to survive treatment with clinical antibiotics and disinfectants by employing a variety of strategies, with persisters and the Eagle effect playing a role in survival to quinolones, tolerance playing a role in survival to ceftriaxone and catalases playing a role in survival to hydrogen peroxide.

## Methods

### Strains and culture conditions

All chemicals were purchased from Merck or Fischer Scientific unless otherwise stated. *Yersinia pseudotuberculosis* strain IP32953 (genome available at https://www.ncbi.nlm.nih.gov/nuccore/BX936398) was kindly provided by the Defence Science and Technology Laboratory. *Y. pseudotuberculosis* was streaked on LB medium agar plates (15 g/L agar, 10 g/L tryptone, 5 g/L yeast extract, and 10 g/L NaCl; Melford) from cryostock. Overnight liquid cultures were prepared by picking a single colony from a streak plate and grown in 200 mL of LB medium and incubated for 17 hours at 37 °C with shaking at 200 rpm.

### Measurement of minimum inhibitory concentration and minimum bactericidal concentration values of antimicrobials against *Y. pseudotuberculosis*

MIC assays were carried out using the 96-well plate microdilution method based on the guidelines set out by the Clinical and Laboratory Standards Institute (CLSI) as previously described (109, 110). Briefly, *Y. pseudotuberculosis* was grown in LB for 17 hours at 37 °C while shaking at 200 rpm. To establish the MIC of antimicrobials against *Y. pseudotuberculosis* in exponential phase, the overnight culture was diluted to an OD_600_ of 0.01 and incubated at 20 °C for a further 6 hours in LB before their addition to the 96-well plate containing antimicrobials at a final bacterial OD_600_ of 0.01.

To determine the MIC of antimicrobials against *Y. pseudotuberculosis* in stationary phase, the overnight culture was diluted in LB and immediately added to the 96-well plate containing antimicrobials at a final bacterial OD_600_ of 0.01. Starting inoculation was calculated at roughly 1×10^6^ CFU. The 96-well plates were incubated for 24 hours at 37 °C while shaking at 200 rpm. After 24 hours the OD_600_ of each individual well was measured and analysed with a plate reader as previously described (111). The average OD_600_ value measured in three wells containing LB only was subtracted from the OD_600_ value in each of all the other wells on the plate to obtain a background subtracted OD_600_ measurement. The MIC value for each antimicrobial in each plate was determined as the minimal antimicrobial concentration for which the associated OD_600_ value was smaller than 10% of the value obtained as an average OD_600_ of untreated wells, i.e. containing *Y. pseudotuberculosis* in LB. All measurements were performed in biological and technical triplicates.

To measure the MBC value of each antimicrobial against *Y. pseudotuberculosis*, the MIC assays were repeated and, after 24 hours, aliquots from wells with antimicrobial concentrations equal or above the MIC value were resuspended in PBS and plated on LB agar plates. Such plates were incubated for 24 hours at 37 °C and the MBC was defined as the lowest antimicrobial concentration required to give a 3 log_10_, i.e. 99.9% killing, reduction in surviving bacteria compared to the initial inoculum (94).

### Time-kill assays

Time-kill assays were performed as previously reported (47, 78). Briefly, an overnight culture of *Y. pseudotuberculosis* was prepared as described above, diluted in 100 ml LB to an OD_600_ of 0.03 (equivalent to around 10^7^ bacteria ml^-1^) and either exposed to each antimicrobial at a concentration of 1×, 10× or 25× its MIC value or incubated in LB only as an untreated control sample. All cultures were incubated at 37°C at 200 rpm for 24 h. Triplicate aliquots were withdrawn from each culture at hourly time points for the first 5 h and after 24 h. Each aliquot was washed in PBS, serially diluted, 10 µL of each dilution were spotted on LB agar plates and incubated at 37 °C, colony-forming units (CFU) were counted on each plate after 24 h incubation. Each experiment was performed in biological triplicate.

The minimum duration of killing (MDK_99_) of each antimicrobial against *Y. pseudotuberculosis* was defined as the minimum duration of treatment that led to a 3 log_10_, i.e. 99.9% killing, reduction in surviving bacteria compared to the initial inoculum (13). Persisters were identified via the observation of biphasic killing with a sharp initial reduction in surviving bacteria followed by a slower reduction and a plateau in the number of survival bacteria; the persister fraction was defined as the ratio between the number of surviving bacteria when this plateau was reached over the number of bacteria in the initial inoculum (15).

For hydrogen peroxide, a second exposure was carried out using cultures already exposed to 2.5, 25, 62.5 and 250 mM hydrogen peroxide for 24 hours. Cultures were resuspended to an OD_600_ of 0.03 with the exception of cultures previously exposed to 250 mM hydrogen peroxide in which the whole culture was centrifuged and resuspended in LB to capture the highest number of cells present. The cultures were resuspended in LB and the same concentration of hydrogen peroxide to which they had been previously exposed. Colony forming unit assays were carried out for each culture as described above.

### Transcriptomic Analysis

RNA extraction, sequencing and analysis were performed on biological triplicate of each condition following previously reported protocols (35, 112). Briefly, overnight cultures of *Y. pseudotuberculosis* were diluted in 100 ml LB to an OD_600_ of 0.03 either with the addition of 2.5 or 62.5 mM of hydrogen peroxide or without addition of hydrogen peroxide. Three 100 µl aliquots were withdrawn from each culture either after 6 h or 24 h transferred into an RNase free tube and centrifuged at 4000 rpm for 15 min. 200 μL of RNA-protect bacteria reagent was added, vortexed for 5 s and then incubated for 5 min at room temperature. Each sample was then centrifuged at 5000 g for 10 min and the supernatant was removed. Extractions were performed using RNeasy minikit (Qiagen) by following the manufacturer’s instructions. DNA removal during extraction was carried out by using RNase-Free DNase I kit (Qiagen). RNA concentration and quality were measured using a Qubit 4.0 fluorometer (Invitrogen^TM^) and 4200 TapeStation (Agilent), respectively, and only samples with an RNA integrity number larger than 7.5 were taken forward and sequenced. Transcript abundance was quantified using Salmon for each gene in all samples. Subsequent differential analysis was performed using DEseq2 in R software to quantify the log_2_ fold change in transcript reads for each gene (113). Clustering analysis and gene ontology enrichment analysis were performed using the mclust package (version 5.4.7) and the clusterProfiler package (version 4.10.0) for R as previously described (80). Enrichment in terms belonging to the ‘Biological Process’ ontology was calculated for each gene cluster, relative to the set of all genes quantified in the experiment, via a one-sided Fisher exact test. p-values were adjusted for false discovery by using the method of Benjamini and Hochberg. The lists of significantly enriched terms were simplified to remove redundant terms, as assessed via their semantic similarity to other enriched terms, using clusterProfiler’s simplify function.

### Statistical analysis

Points and error bars displayed in all graphs represent the mean and standard error of the mean. Biological triplicate experiments were conducted in all instances. All graphs were generated using GraphPad Prism 9.

## Acknowledgments

This work was supported by the BBSRC, the EPSRC, and the MRC through three grants awarded to S.P. (BB/V008021/1, EP/Y023528/1, and MR/Y033892/1). O.L., C.H.J. and I.H.N. were supported by the Defence Science and Technology Laboratory. This project also utilised equipment funded by a Wellcome Trust Institutional Strategic Support Fund (WT097835MF), a Wellcome Trust Multi-User Equipment Award (WT101650MA) and a BBSRC award (BB/Z515942/1).

The funders had no role in study design, data collection and analysis, decision to publish, or preparation of the manuscript. For the purpose of open access, the authors have applied a Creative Commons Attribution (CC BY) licence to any Author Accepted Manuscript version arising from this submission.

## Author contributions

Conceptualization: S.P.; methodology: O.G., J.W., U.L.; formal analysis: O.G., J.W., U.L., J.O., A.F., A.J., C.H.J., I.H.N., S.P.; generation of figures: O.G., J.W., U.L. and S.P.; investigation: O.G., J.W., U.L., J.O., A.F., A.J., C.H.J., I.H.N., S.P.; resources: S.P.; data curation: O.G., J.W., U.L., J.O., A.F., A.J., C.H.J., I.H.N., S.P.; writing - original draft: O.G., J.W., S.P.; writing – review & editing: O.G., J.W., U.L., J.O., A.F., A.J., C.H.J., I.H.N., S.P.; visualisation: O.G., J.W., U.L. and S.P.; supervision: I.H.N., S.P.; project administration: S.P.; funding acquisition: I.H.N., S.P.

## Data availability

All data generated or analyzed during this study are included in this published article and its supplemental files.

## Supplementary Material

**Supplementary Figure 1.**
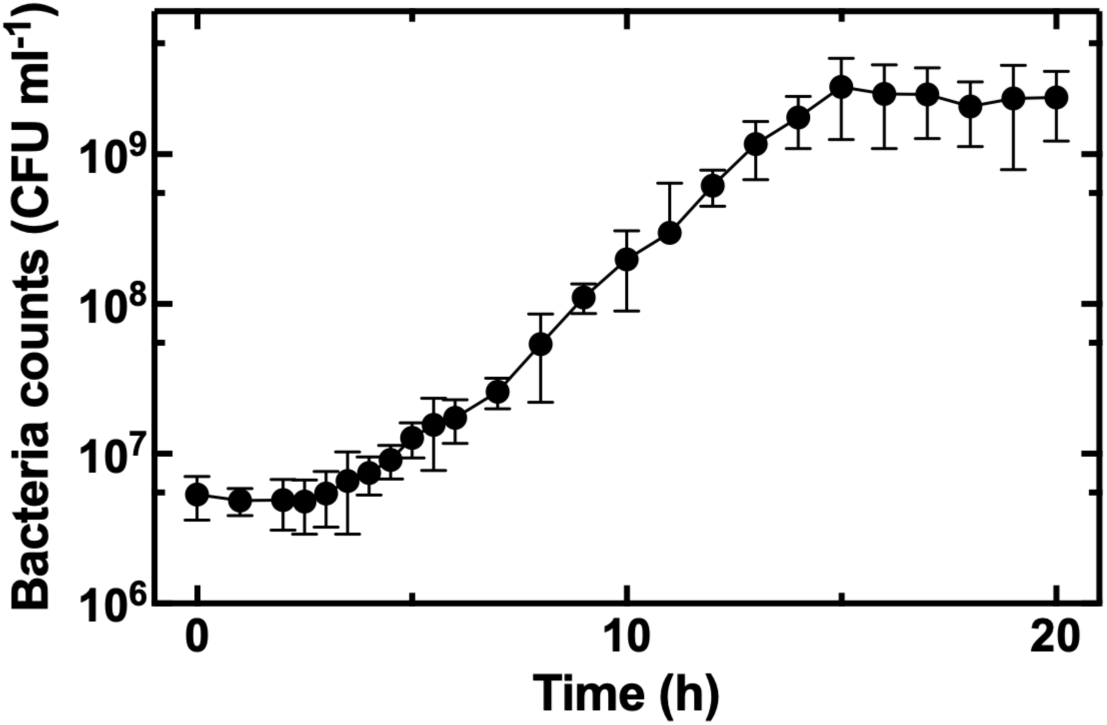
Temporal dependence of bacteria counts for *Y. pseudotuberculosis* growing in LB medium at 37 °C. The data points and error bars were calculated as the mean and standard deviation of CFU measurements carried out in biological triplicate each consisting of technical triplicate.

**Supplementary Figure 2.**
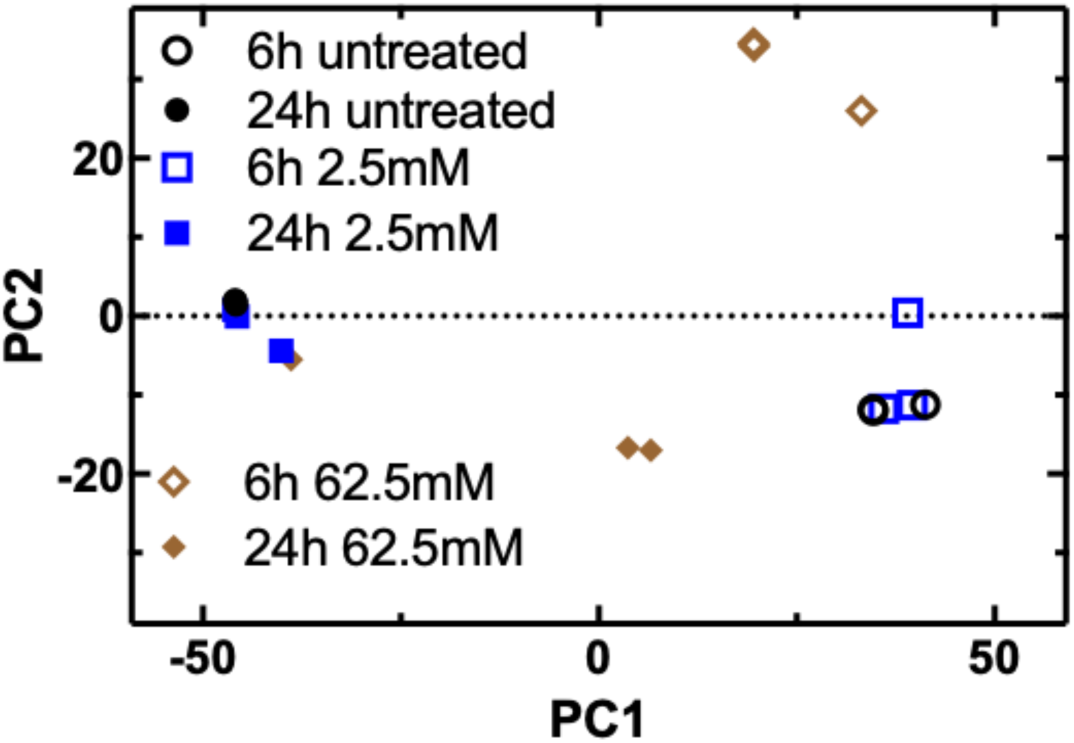
Principal component analysis of replicate transcriptomes of *Y. pseudotubercolosis* after 6 h (open symbols) or 24 h (filled symbols) of incubation in LB (circles) or exposure to 2.5 mM (squares) or 62.5 mM hydrogen peroxide (diamonds). Each transcriptome was measured in biological triplicate.

**S1 File** Transcript counts for each aligned *Y. pseudotubercolosis* gene and for each replicate of each experimental condition, i.e. after 6 h or 24 h of either incubation in LB or exposure to 2.5 mM or 62.5 mM hydrogen peroxide.

**S2 File** Differential expression of *Y. pseudotubercolosis* genes after 6 h or 24 h of exposure to 2.5 mM or 62.5 mM hydrogen peroxide compared to 6 h or 24 h incubation in LB. Gene number, cluster number, gene name, gene ontology (go) term, go function, product, go process, transcript base mean, log_2_ fold change, standard error of log_2_ fold change, p-value, adjusted p-value, gene identifier are reported for each *Y. pseudotubercolosis* gene and each comparison.

**S3-S10 Files** Gene ontology enrichment analysis of differentially expressed *Y. pseudotubercolosis* in each cluster identified via cluster analysis. GO Identifier, description, relative number of genes contained within the cluster and the genome, p-value, adjusted p-value, q-value and identifier of each gene are reported for each significantly enriched biological process.

